# Enhanced neutralization potency of an identical HIV neutralizing antibody expressed as different isotypes is achieved through genetically distinct mechanisms

**DOI:** 10.1101/2022.08.17.504233

**Authors:** Thandeka Moyo-Gwete, Cathrine Scheepers, Zanele Makhado, Prudence Kgagudi, Nonkululeko B. Mzindle, Rutendo Ziki, Sharon Madzorera, Nelia P. Manamela, Frances Ayres, Bronwen E. Lambson, Simone I. Richardson, Lynn Morris, Penny L. Moore

## Abstract

Antibodies with the same variable region can exist as multiple isotypes with varying neutralization potencies, though the mechanism for this is not fully defined. We previously isolated an HIV-directed IgA1 monoclonal antibody (mAb), CAP88-CH06, and showed that IgA1 and IgG3 isotypes of this antibody demonstrated enhanced neutralization compared to IgG1. To explore the mechanism behind this, hinge region and constant heavy chain (CH1) chimeras were constructed between the IgA1, IgG3 and IgG1 mAbs and assessed for neutralization activity, antibody-dependent cellular phagocytosis (ADCP) and antibody-dependent cellular cytotoxicity (ADCC). Hinge chimeras revealed that the increased neutralization potency and phagocytosis of the IgG3 isotype was attributed to its longer hinge region. In contrast, for IgA1, CH1 chimeras showed that this region was responsible both for enhanced neutralization potency and decreased ADCP, though ADCC was not affected. Overall, these data show that the enhanced neutralization potency of CAP88-CH06 IgG3 and IgA1, compared to IgG1, is achieved through distinct mechanisms. Understanding the influence of the hinge and CH1 regions on Fab domain function may provide insights into the engineering of therapeutic antibodies with increased neutralization potency.

## Introduction

Antibodies are B-cell-derived Y-shaped proteins capable of neutralizing pathogens through the antigen-binding Fab region and recruiting innate immune cell functions through Fc binding to a Fc receptor (FcR) [1]. The Fab is comprised of the variable heavy and light chain regions, which form the antibody paratope that binds its target, as well as the constant heavy chain 1 (CH1) and constant light chain (C_L_) regions [1]. The structure of the Fc region varies by isotype, and this is a major modulator of Fc effector function. While the Fab is primarily responsible for antigen binding, several studies have shown that the constant region also alters the binding affinity and neutralization of antibodies against a variety of antigens and pathogens [2, 3]. Although the majority of therapeutic antibodies on the market are expressed as IgG1, other isotypes may be valuable to explore as their unique structural characteristics may enhance the polyfunctionality of the mAbs [4, 5].

There is significant sequence variation in both the hinge and CH1 regions that differentiates the constant regions of various isotypes and both these regions have been implicated in influencing neutralization potency [6, 7]. The increased neutralization potency of an IgG3 version of the HIV-specific V2-apex-specific broadly neutralizing antibody (bNAb), CAP256-VRC26.25, was attributed to the longer hinge region [6]. In contrast, the IgA2 CH1 region of the anti-HIV bNAb 2F5 improved binding affinity to the membrane proximal external region (MPER) [7]. However, whether these mechanisms are used by antibodies targeting different HIV epitopes has not been explored [8].

We previously isolated an IgA1 monoclonal antibody (mAb), CAP88-CH06, from an HIV-infected donor, CAP88, that targets the C3/V4 region and potently neutralizes early autologous viruses [9]. Although very potent, this antibody is strain-specific and does not neutralize heterologous viruses. While exploring the role of class-switching within this lineage, we found co-circulating antibodies representing multiple isotypes including IgG1 and IgG3 – some of which shared identical variable regions, but exhibited markedly different neutralization activity [10]. Furthermore, when the CAP88-CH06 antibody was engineered as an IgG1, it exhibited reduced potency against autologous viruses, while an IgG3 engineered form of the antibody displayed high potency, comparable to the original IgA1 isotype [9].

Therefore, to define the mechanism for increased neutralizing potency in IgA1 and IgG3 isotypes, we generated antibody chimeras, separately swapping the hinge regions and the CH1 regions between the IgA1, IgG3 and IgG1 isotypes. Our data show that the extended hinge region of the CAP88-CH06 IgG3 mAb drives the enhanced neutralization potency of this isotype, whereas the CH1 region drives that of the IgA1 isotype. This indicates that there are distinct genetic mechanisms for achieving enhanced breadth, with implications for the engineering of enhanced mAbs.

## Results

### The IgG3 hinge region enhances CAP88-CH06 neutralization potency

The naturally isolated CAP88-CH06 IgA1 and engineered IgG3 and IgG1 mAbs all share identical Fab variable regions but differ in their constant regions. To determine their neutralization potency, mAbs were tested against 12 early autologous viruses isolated from donor CAP88 between 5-30 weeks post-infection. As we previously showed [10], the IgG1 isotype displayed the least potency (geometric mean titer (GMT): 0.30) followed by the IgG3 isotype (GMT: 0.06). The CAP88-CH06 IgA1 mAb was the most potent of the three (GMT: 0.02) (**Fig. 1A**).

**Figure 1:**
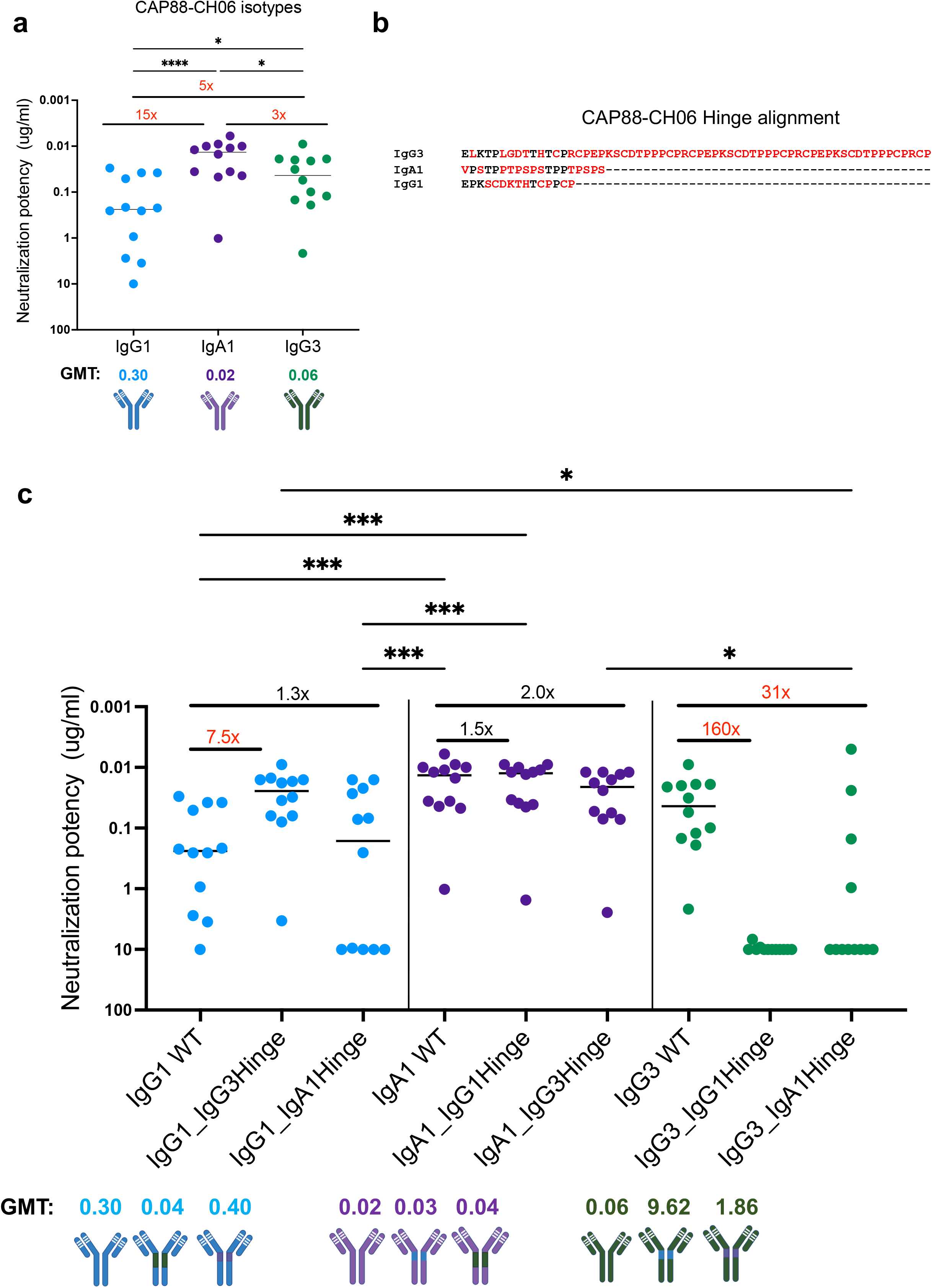
The extended IgG3 hinge results in increased neutralization potency in CAP88-CH06 antibodies. (**a**) CAP88-CH06 IgG3 (green) and IgA1 (purple) and IgG1 (blue) mAbs when tested for neutralization against 12 CAP88 autologous viruses in a pseudotyped virus neutralization assay. (**b**) Alignment of the CAP88-CH06 mAb hinge regions performed using Aliview v1.20. Red denotes amino acids that differ between hinge sequences mAbs and dashes represent gaps. (**c**) Antibody hinge chimeras were tested for neutralization against 12 CAP88 autologous viruses. All experiments were performed in duplicate with at least two replicates. Neutralization potency is shown as IC_50_ values in ug/ml with neutralization resistance being defined as >10 ug/ml. GMT: geometric mean titre. Fold changes are shown in red (if greater than 3-fold different) or black (if less than 3-fold different). Stasticical significance was calculated using the Friedman test with Dunns correction for multiple comparisons. Significance is shown as: *p<0.05, **p<0.01, ***p<0.001 and ****p<0.0001. Antibody schematics were created in Biorender (www.biorender.com).

We first made chimeric versions of these three mAbs with the hinge regions swapped between them. We postulated that the varying lengths, sequences and overall structure of the hinges may play a role in the flexibility of the antibody and access to its epitope, thereby impacting neutralization. The hinge of IgG3 (IgG3*01) is 62 residues long while IgG1 (IgG1 *01) has a hinge of 15 residues and IgA1 (IgA1 *01) of 19 residues (**Fig. 1B**, with residues that vary between hinges shown in red). The WT and the chimeras all bound to an autologous CAP88 gp120 protein, with binding not being significantly different between the antibodies (**Supplementary Figure 1**).

When neutralization was assessed, we found that the IgG1 antibody was 7.5-fold more potent when it contained the extended hinge of the IgG3 antibody (IgG1_IgG3Hinge GMT: 0.04) (**Fig.1C**). Introduction of the IgA1 hinge region however did not significantly affect potency of the IgG1 mAb (1.3-fold increase in GMT) (**Fig.1C**). Engineering of the IgG1 hinge into the IgA1 mAb had no impact on the neutralization potency (GMT: 0.02 vs 0.03). However, substitution of the IgG3 hinge into the IgA1 mAb resulted in a 2-fold decrease in potency for the IgA1 mAb, consistent with the fact that the potency of the IgG3 WT was lower than that of the IgA1 WT (WT GMT: 0.06 and 0.02, respectively) (**Fig. 1A**). The GMT of the IgA1_IgG3Hinge chimera was similar to the GMT of the IgG3 WT (GMT: 0.04 vs 0.06). The exchange of either an IgG1 or an IgA1 hinge to the IgG3 mAb, resulted in a significant reduction in neutralization potency of the IgG3 mAb (160-fold, GMT of 9.62 for IgG3_IgG1Hinge and 31-fold, GMT of 1.86 for IgG3_IgA1Hinge) (**Fig. 1C**). Together, these data indicate that the hinge region is a modulator of neutralization potency for the CAP88-CH06 IgG3 mAb, but not for the IgA1 mAb.

As hinge length is also known to modulate Fc effector function [6, 11], we assessed the ability of the hinge chimeras to mediate antibody dependent cellular phagocytosis (ADCP) and antibody dependent cellular cytotoxicity (ADCC). Using an CAP88 autologous virus from 30 weeks post infection we showed that both the IgG3 and IgG1 parental mAbs had strong phagocytosis activity. When extended this to a subset of the chimeric mAbs, we found that ADCP mediated by the IgG1 mAb trended towards slightly increased activity by the addition of the IgG3 hinge (IgG1_IgG3Hinge, 1.3x-fold higher), to levels similar to the IgG3 WT (**Fig. 2A**). Substitution of the IgG3 hinge with that of IgG1 (IgG3_IgG1Hinge) resulted in reduced ADCP relative to the parental IgG3 antibody. For ADCC, the IgG3 and IgG1 WT mAbs showed similar levels of activity, but the IgG3 antibody which contained the IgG1 hinge had reduced ADCC (**Fig. 2B**). Altogether, our findings suggest that the IgG3 hinge results in both increased neutralization and ADCP, with no improvement in ADCC.

**Figure 2:**
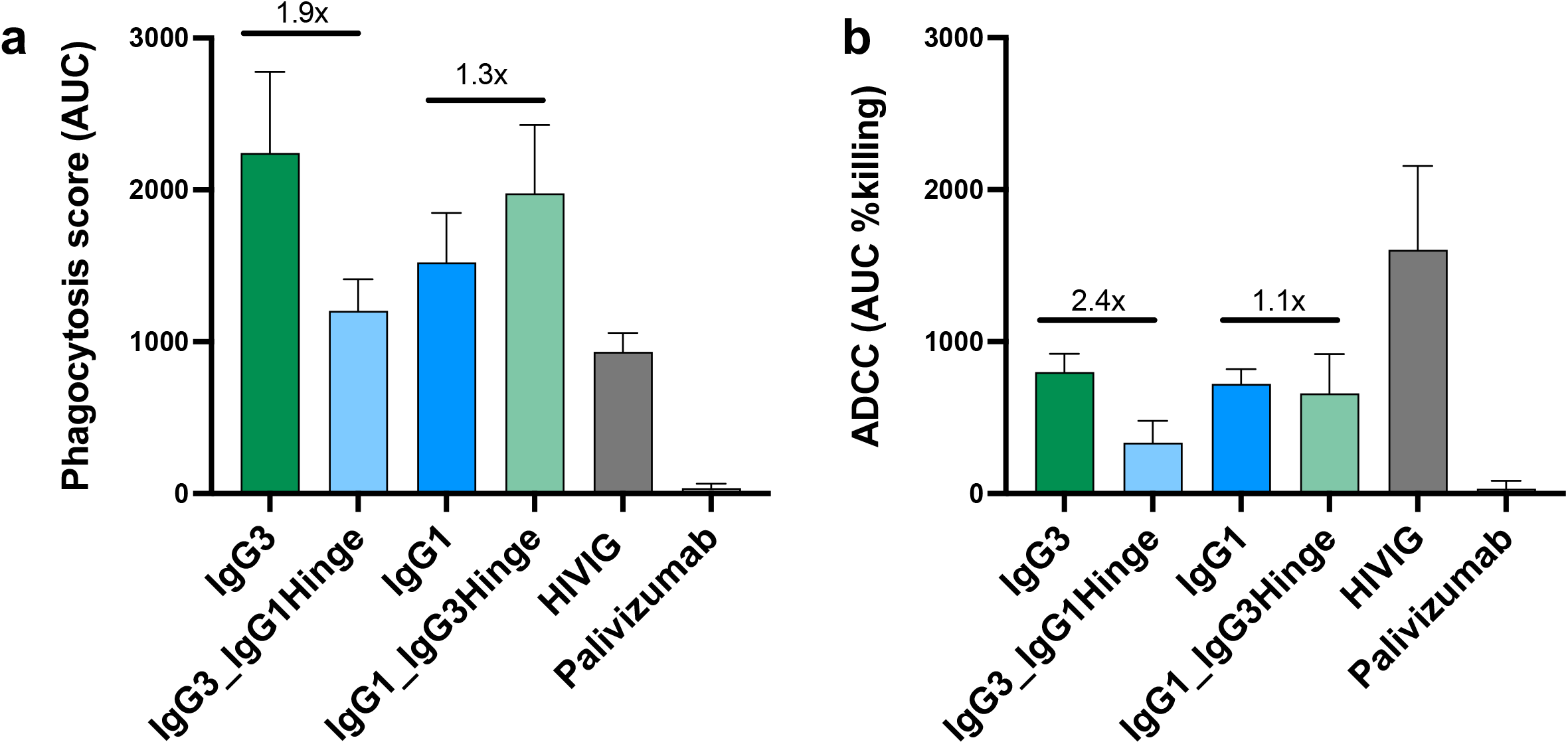
The IgG3 hinge results in a miminal increase in ADCP and does not substantially affect ADCC in CAP88-CH06 antibodies. (**a**) Antibody-dependent cellular phagocytosis (ADCP) activity of the CAP88-CH06 IgG3 WT, IgG3_IgG1Hinge, IgG1 WT and IgG1_IgG3Hinge antibodies was measured using THP-1 phagocytosis assay and the phagocytosis score shown as an area under the curve (AUC) measure. (**b**) Antibody dependent cellular cytotoxicity (ADCC) activity of the CAP88-CH06 IgG3 WT, IgG3_IgG1Hinge, IgG1 WT and IgG1_IgG3Hinge antibodies was measured using an infectious ADCC assay. The % killing activity is shown as area under the curve (AUC) measure. In all experiments, HIVIG, a polyclonal plasma cocktail from HIV-infected individuals, was used as a positive control and Palivizumab, a RSV-specific mAb, was used as a negative control. Experiments were conducted in duplicate and error bars represent the mean with standard deviation of two experiments. The Krustal-Wallis test with Dunn’s correction was performed and no comparisons were significantly different, except for the negative control, Palivizumab. Fold changes are shown in black.

### The IgA1 CH1 region contributes to increased CAP88-CH06 neutralization potency

As the CH1 region has been implicated in increased potency of an IgA2 mAb [7], we investigated the role of this region in the increased neutralization potency of the CAP88 IgA1 isotype. The IgG1 and IgG3 CH1 regions differ by 4 residues (shown in red) whereas the IgA1 CH1 differs by more than 60% compared to that of the IgG1 and IgG3 mAbs, including four insertions [12] (**Fig. 3A**). We swapped the CH1 regions between the three mAbs and tested them for neutralization using the same 12 viruses tested above (**Fig. 3B**). As with the hinge swap chimeras, all the chimeras were able to bind to a CAP88 autologous gp120 protein and no significant differences were observed in binding ability (**Supplementary Figure 1**).

**Figure 3:**
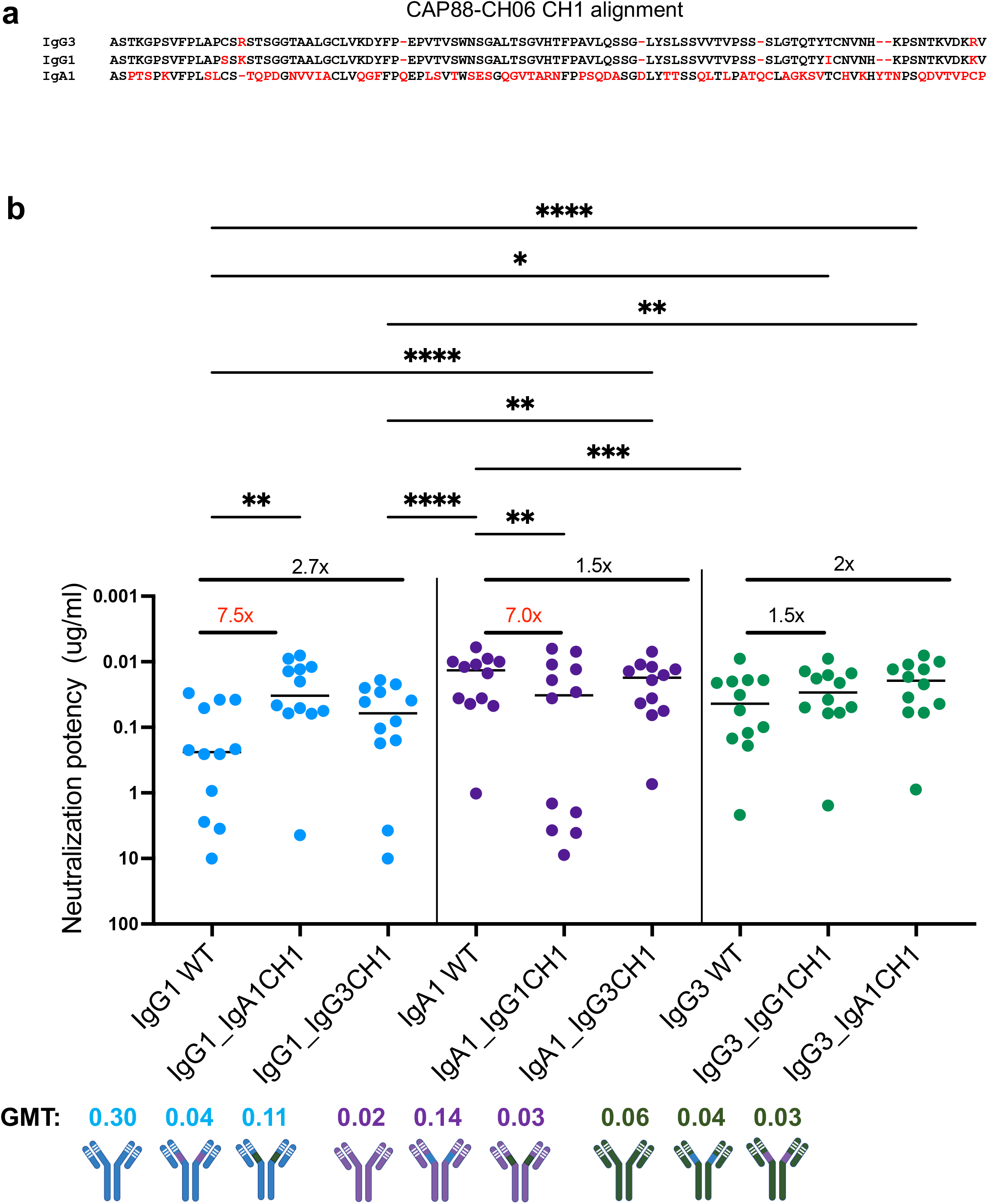
The IgA1 CH1 region results in increased neutralization potency in CAP88-CH06 antibodies. (**a**) Alignment of CAP88-CH06 mAb CH1 regions performed with Aliview v1.20. Red denotes amino acid differences between the CH1 sequences of the three mAbs. (**b**) Antibody CH1 chimeras were tested for neutralization against 12 CAP88 autologous viruses. All experiments were performed in duplicate with at least two replicates. Neutralization potency is showns as IC_50_ values in ug/ml with neutralization resistance being >10 ug/ml. GMT: geometric mean titre. Fold changes are shown in red (greater than 3-fold different) or black (less than 3-fold different). Stasticical significance was calculated using the Friedman test with Dunns correction for multiple comparisons. Significance is shown as: *p<0.05, **p<0.01, ***p<0.001 and ****p<0.0001. Antibody schematics were created in Biorender (www.biorender.com).

The IgG1 antibody increased in potency by 7.5-fold from a GMT of 0.30 to 0.04 upon replacement with the IgA1 CH1 (IgG1_IgA1CH1), resulting in potency similar to the IgA1 WT (GMT: 0.02) (**Fig. 3B**). The exchange with the IgG3 CH1 (IgG1_IgG3CH1) slightly increased the potency of the IgG1 mAb (from GMT 0.30 to 0.11) (**Fig. 3B**). Removal of the IgA1 CH1 from the IgA1 WT and replacement with the IgG1 CH1 (IgA1_IgG1CH1) resulted in a decrease in neutralization potency (from GMT: 0.02 to 0.14) whilst replacement of the IgG3 CH1 with the IgA1 mAb (IgA1_IgG3CH1) did not result in a significant change in potency (GMT: 0.02 vs 0.03) (**Fig. 3C**). Although the exchange of the IgG1 CH1 into the IgG3 mAb (IgG3_IgG1CH1) resulted in a change to the potency of the IgG3 mAb (from GMT: 0.06 to 0.04), a slighty higher increase in potency was observed when the IgA1 CH1 was swapped into the IgG3 mAb with IgG3_IgA1CH1 mAb nearly recapitulating the potency of the IgA1 WT mAb (GMT: 0.06 to 0.03; IgA WT GMT: 0.02) (**Fig. 3D**).

We next assessed the role of the CH1 on ADCP and ADCC (**Fig. 4**). In ADCP assays, the IgA1 mAb had a trend towards a reduced phagocytic score upon addition of the IgG1 CH1 region (1.8-fold drop) (**Fig. 4B**). However, insertion of the IgA CH1 into IgG1 did not confer improved ADCP but in fact resulted in considerable loss of ADCP activity. For ADCC, the effect of swapping CH1 regions between IgA1 and IgG1, compared to the parental antibodies was minor (1.4- and 1.6-fold, respectively), likely indicating that other structural elements of the mAb mediate this function. Overall, these results strongly suggest that the CAP88-CH06 IgA1 CH1 drives the increased neutralization potency of this mAb but does not significantly impact ADCC, and reduces ADCP.

**Figure 4:**
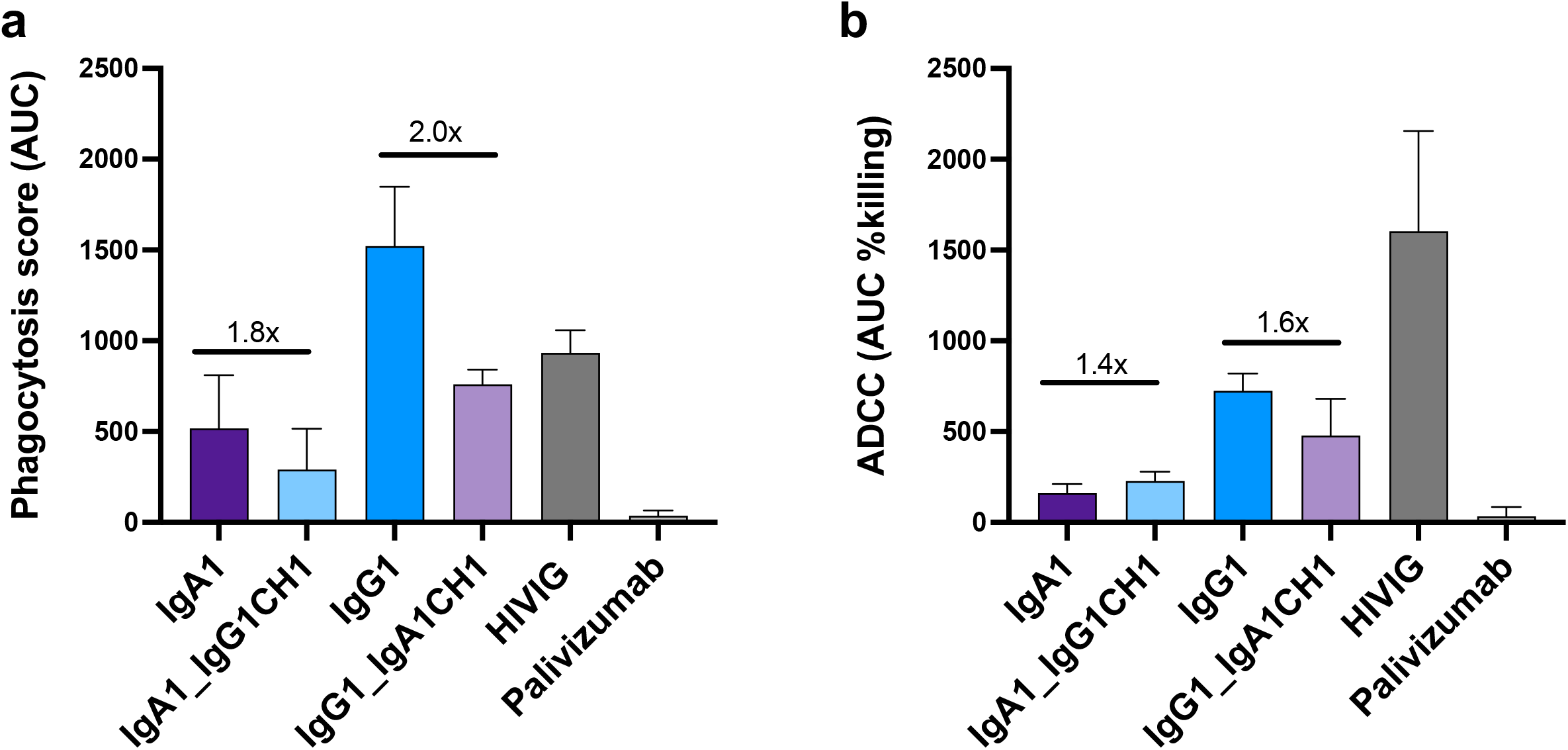
The IgA1 CH1 region decreases ADCP and has little effect on ADCC in CAP88-CH06 antibodies. (**a**) ADCP activity of the CAP88-CH06 IgA1 WT, IgA1_IgG1CH1 and IgG1_IgA1CH1 antibodies was measured using THP-1 phagocytosis assay and the phagocytosis score shown as an area under the curve (AUC) measure. (**b**) ADCC activity of the CAP88-CH06 IgA1 WT, IgA1_IgG1CH1, IgG1 WT and IgG1_IgA1CH1 antibodies was measured using an infectious ADCC assay. The % killing activity is shown as an AUC measure. In all experiments, HIVIG, a polyclonal plasma cocktail from HIV-infected individuals, is used as a positive control and Palivizumab, a RSV-specific mAb, was used as a negative control. Experiments were conducted in duplicate and error bars represent the mean with standard deviation of two experiments. The Krustal-Wallis test with Dunn’s correction was performed and no comparisons were significantly different except for the negative control, Palivizumab. Fold changes are shown in black.

## Discussion

While the role of isotype in Fc effector function is well known, the impact of the Fc on neutralization and binding capacity of antibodies has only recently gained attention [6, 7, 10, 13, 14]. We previously isolated a neutralizing antibody, CAP88-CH06 and studied the maturation of this lineage identifying co-circulating antibodies of different isotypes with identical Fab regions [9, 10]. Furthermore, we found that certain class-switching events resulted in increased neutralization potency conferring the ability of isotypes to tolerate certain antibody escape mutations [10]. To probe the mechanism behind this, we explored the differences in neutralization between the naturally-occuring IgA1 CAP88-CH06 and engineered IgG1 and IgG3 versions of CAP88-CH06, all with identical antigen-binding regions. We show that the neutralization potency of the CAP88-CH06 IgG3 isotype is mediated by its long hinge region while the increased neutralization capacity of the IgA1 mAb is modulated by the CH1 region of the antibody. These data show that there are multiple ways to achieve increased neutralization within a single antibody that developed in an HIV-infected individual.

Varied neutralization potency mediated by the Fc region is not unique to the CAP88-CH06 antibody, with several other studies reporting the influence of the Fc region on neutralization and binding on anti-HIV-1 antibodies. The IgG3 isotypes of several antibodies have been shown to be more potent than their IgG1 versions, including CAP256-VRC26.25, 35O22 and PGT 135, though this effect was virus-specific [8]. Cheeseman *et al.*, found that the IgA versions of some anti-HIV-1 antibodies they tested were less or completely ineffective at inhibiting HIV-1 infection compared to their IgG counterparts [15]. Similarly, when the CD4 binding site-directed antibody CH31 was expressed as an IgG1, it had higher neutralization activity and was more protective than its IgA2 version in mucosal infection models [16]. This effect is likely epitope- or assay-specific as the IgA2 version of the MPER-specific 2F5 mAb displayed improved affinity and neutralization activity by compared to IgG [7]. Further studies are needed to establish how generaliseable these finding.

The hinge region varies markedly by isotype, and is substantially longer in IgG3 antibodies. We have previously shown that the hinge region in IgG3 antibodies can enhance polyfunctionality of an HIV-directed, anti-V2 apex antibody (Richardson et al., 2019) but this is the first report on the importance of the IgG3 hinge region for a C3/V4-targeting, glycan-dependent HIV-specific antibody. The increased length of the IgG3 hinge likely increases the flexibility of the Fab region [17] potentially providing greater ability to maneuver past the HIV glycan shield and access its epitope. In addition, the sequence of the IgG3 hinge is very different to those of IgG1 and IgA1, which likely results in a distinct structural conformation leading to differential affinity of the IgG3 Fab to antigens [18]. The Fab-Fab distance in IgG3 antibodies is far greater than other Ig antibodies which may further influence binding affinity and neutralization potential [18–20]. In addition to the long IgG3 hinge length and increased Fab-Fab angles, this region also possesses O-linked glycosylation sites [21]. As the role of O-linked glycosylation on antibody function, binding and neutralization has not been extensively studied [22], further investigation into the role of this type of glycosylation should be conducted to assess whether this also impacts antibody neutralization. In this study, all antibodies were produced in human embryonic kidney (HEK) 293F cells. However, expression of the antibodies in cell lines which bias the expression of certain glycan forms and other post-translational modifications, allied with approaches such as mass spectrometry to confirm the exact types of glycans present would be essential for future work.

The CAP88-CH06 IgA1 antibody uses a distinct mechanism to achieve increased neutralization compared to the IgG3 isotype. The IgA1 CH1 is 60% different at the amino acid level, compared to both IgG1 and IgG3 isotypes. There is an additional deletion and a number of insertions, all of which likely change the conformation of the IgA1 CH1 region. These differences may result in increased affinity of interactions between the IgA1 CH1 and variable regions which may play a role in influencing Fab interactions [12, 23]. The IgA1 CH1 region has increased flexibility which has been shown to influence paratope-epitope binding kinetics [7]. This is one of the few reports implicating the CH1 region in enhancing neutralization capacity of an antibody and requires further study, particularly through atomic level structural analysis. However, our study also highlights the fact that construction of chimeras can also result in loss of function. While introduction of the IgA1 CH1 region into IgG1 was associated with increased neutralization, we also observed reduced ADCP in this antibody, as well as the reverse chimera. These sometimes unpredictable functional effects highlight the complexity of engineering both potency and polyfunctionality into antibodies.

Our study did not investigate how viral signatures may have contributed to the increase in potency of the IgG3 and IgA1 mAbs. When we tested the binding ability of the chimeras to determine if the differences observed in neutralization would correlate with binding avidity, no significant differences were observed between the WTs and chimeras, implicating other factors in modulating neutralization. We have previously shown that variable neutralization potency of IgG3 allelic variants of mAb CAP256.25 was dependent on the presence or absence of viral glycans within the epitope [6]. Within the panel of CAP88 viruses tested, the effect on neutralization potency of chimeric antibodies varied. For example, while introduction of the IgA1 hinge into IgG1 resulted in an overall 1.3 fold increase in GMT, several viruses were resistant to the chimeric antibody. This observation suggests a role for viral signatures and epitope differences in influencing how big an effect the hinge and CH1 regions have on the potency of the CAP88 mAbs, and warrants further study.

The use of monoclonal antibodies for therapeutic and prophylactic use has dramatically increased in recent years [24, 25]. The development of potent therapeutic mAbs with longer half-lives is a challenge in the field, as antibodies are costly, and therefore novel approaches to the engineering of these mAbs are being investigated. These data suggest that the IgG1 isotype, which has a longer half-life than both IgG3 and IgA1 isotypes [26, 27], could be engineered for greater neutralization by incorporating an IgG3 hinge region or an IgA1 CH1 region. The multiple mechanisms in which antibodies can achieve increased function extends the options for engineering enhanced antibodies while retaining half-lives. Furthermore, there is considerable functionally unexplored genetic diversity within antibody genes/allotypes [6, 28–30] and expanding mechanistic studies to incorporate this diversity may further enhance engineering efforts.

The differential ability of isotypes to neutralize pathogens also has implications for active vaccination. We previously showed that class-switching within the CAP88-CH06 lineage enabled continuing affinity maturation despite viral escape from some isotypes [10]. This continued maturation, which is a function of the varying ability of different isotypes to neutralize viruses with different mutations, suggests the benefit of enhancing class-switching in vaccine regimes. Indeed we have previously shown that HIV infected individuals who develop broadly neutralizing antibodies have increased IgG subclass diversity during early infection [31].-Together these data suggest possibilities to manipulate the natural landscape of class-switching to increase antibody potency and possibly breadth. Adjuvants which promote class-switching may be beneficial in inducing potent neutralizing responses in vaccinees and should be investigated further in the context of an HIV vaccine [32–35].

Overall, our data shed light on the key regions used by the constant region of an antibody to influence neutralization of the HIV Env. These data have implications for design of therapeutic mAbs and for adjuvanting of vaccines, where elicitation of multiple isotypes is likely to contribute to high levels of polyfunctionality. As most HIV-1 mAbs are expressed as IgG1 isotypes, exploring the use of IgA1 or IgG3 mAbs and further understanding how the CH1 and hinge regions in other isotypes influence neutralization and function may increase the potency of therapeutic mAbs for HIV-1 and other pathogens.

## Materials and Methods

### Design and expression of CAP88-CH06 Hinge and CH1 swap chimeras

The CAP88-CH06 variable region was cloned into IgG1, IgA1 and IgG3 CMVR backbones using the NEBuilder® HiFi DNA Assembly Cloning Kit. Sanger sequencing was used to assess the success of the cloning reactions. Human embryonic kidney (HEK) 293F suspension cells were transfected with heavy and light chain plasmids using the PEImax transfection reagent. After incubating for six or seven days at 37°C, 70% humidity and 10% CO_2_, proteins were purified using a Protein A resin, followed by buffer exchange into 1x PBS buffer and freezing at −80 °C until use.

### HIV-1 pseudovirus-based neutralization assay

The pseudovirus-based neutralization assay was performed as previously described [36]. Briefly, monoclonal antibodies were incubated for 60-90 minutes with HIV pseudovirus which was made by co-transfecting the HEK 293T cell line with an HIV-1 Env plasmid of interest and a pSG3 backbone. After incubation, TZM-bl cells were added followed by a further incubation of 48 hours upon which the assay readout was measured by the level of luminescence. An inhibitory concentration where 50% of the virus is neutralized (IC_50_) was calculated and graphs plotted in Graphpad Prism 9.0.1.

### HIV-1 gp120 enzyme linked immunosorbent assay

96-well high binding plates were coated with 2 μg/ml of CAP88 3.10.29 gp120 core protein and incubated overnight at 4 ^0^C. After washing with 1x PBS + 0.05% Tween 20 and blocking with a buffer containing 1x PBS, 0.05% Tween 20 and 5% milk powder, the CAP88 monoclonal antibodies were added in a 3-fold dilution series from a starting concentration of 1 ug/ml. After a 1 hour incubation at 37 ^0^C, a horseradish peroxidase-conjugated secondary antibody diluted at 1:3000 was added followed by a subsequent incubation step at 37 ^0^C for 1 hour. 3,3’,5,5’-tetramethylbenzidine (TMB) substrate was added and after 5 minutes the reaction was stopped using 1M sulfuric acid. The level of binding was measured at an optical density of 450 nm and analysis was conducted in Graphpad Prism v9.4.1.

### Antibody-dependent cellular phagocytosis assay (ADCP)

The THP-1 phagocytosis assay was conducted as previously described [37] using 1μM neutravidin beads coated with an autologous CAP88 gp120 protein (CAP88 3.10.29). Monoclonal antibodies were tested starting at 50 μg/ml with 5-fold dilutions. Phagocytic scores were calculated as the geometric mean FITC-A fluorescence that the beads that have taken up multiplied by the percentage bead uptake. Pooled IgG from HIV-positive donors (HIVIG) (NIH AIDS Reagent Program) was used in all assays to normalize the data. Palivizumab was used as a negative control.

### Antibody-dependent cellular cytotoxicity (ADCC)

Experiments were conducted using replication-competent infectious molecular clone (IMC) encoding the CAP88 3.10.29 *env* within an isogenic backbone Env-IMC-6ATRi, as described previously [38]. These constructs were kindly provided by Dr Christina Ochsenbauer (University of Alabama at Birmingham). Reporter virus stocks were generated by transfection of HEK 293T cells with proviral IMC plasmid DNA, and titered for infectivity in CEM.NKRCCR5 cells (NIH AIDS Reagent Program) by p24 staining (Beckman-Coulter). ADCC activity was measured as previously described [39]. Briefly, the CEM.NKRCCR5 cell line was used as the target of infection with the IMC. These were incubated with mAbs starting at 50 μg/ml. Cryopreserved peripheral blood mononuclear cells (PBMC) obtained from a HIV-negative donor were used as source of effector cells. The cryopreserved PBMCs were rested overnight and effector cells, target cells, and Ab dilutions were plated in white 96-well half area plates and incubated for 6 hours at 37°C in 5% CO_2_. The final readout was the luminescence intensity generated by the presence of residual intact target cells that had not been lysed by the effector population in the presence of any ADCC-mediating mAb. Pooled IgG from HIV-positive donors (HIVIG) (NIH AIDS Reagent Program) was used in all assays to normalize the data. Palivizumab (Medimmune; Synagis) and A32 (NIH AIDS Reagent Program) were used as negative controls.

## Supporting information

Supplementary Figure 1: CAP88-CH06 antibodies and their hinge and CH1 chimeras bind to the HIV-1 gp120

## Acknowledgments

We acknowledge research funding from the South African Medical Research Council (SAMRC) SHIP program and the Centre for the AIDS Program of Research (CAPRISA). CAPRISA is funded by the South African HIV/AIDS Research and Innovation Platform of the South African Department of Science and Technology and was initially supported by the U.S. NIAID, NIH, U.S. Department of Health and Human Services grant U19 AI51794. P.L.M. is supported by the South African Research Chairs Initiative of the Department of Science and Innovation and the National Research Foundation of South Africa (grant no. 98341).

## Author contributions

TMG, CS, LM and PLM designed the study. TMG analyzed the data. TMG and PLM wrote the manuscript. TMG, ZM, FA and BEL designed and expressed the antibody constructs. PK, NBM, RZ and SM performed neutralization assays. NPM and SIR performed ADCP and ADCC assays and analyzed the data. All authors reviewed and edited the manuscript.

## Data availability

The heavy and light chain antibody variable region sequences used to generate the CAP88-CH06 antibodies were deposited into the Genbank database with heavy chain accession number: MN228652.1 and light chain accession number: MN228657.1. The viruses used in this study have been deposited into GenBank and their accession numbers are: MK206069.1, MK206156.1, MK206116.1, MK205479.1, MK206074.1, MK206112.1, MK205486.1, MK205490.1, MK205487.1, MK205495.1.

---

**Supplementary Figure 1: CAP88-CH06 antibodies and their hinge and CH1 chimeras bind to the HIV-1 gp120.** The CAP88-CH06 (**a**) IgG1, (**b**) IgA1 and (**c**) IgG3 WT antibodies and their respective hinge and CH1 chimeras were tested for binding to an autologous CAP88 gp120 protein using an in-house ELISA. All experiments were conducted in duplicate and error bars represent the mean with standard deviation of three experiments. The area under the curve of each antibody titration curve was calculated and the Krustal-Wallis test with Dunn’s correction was performed. No comparisons were significantly different.

