## Supplementary figures and images for "Enhanced neutralization potency of an identical HIV neutralizing antibody expressed as different isotypes is achieved through genetically distinct mechanisms"

### Supplementary Figure 1: CAP88-CH06 antibodies and their hinge and CH1 chimeras bind to the HIV-1 gp120

#### Antibody binding to CAP88 gp120

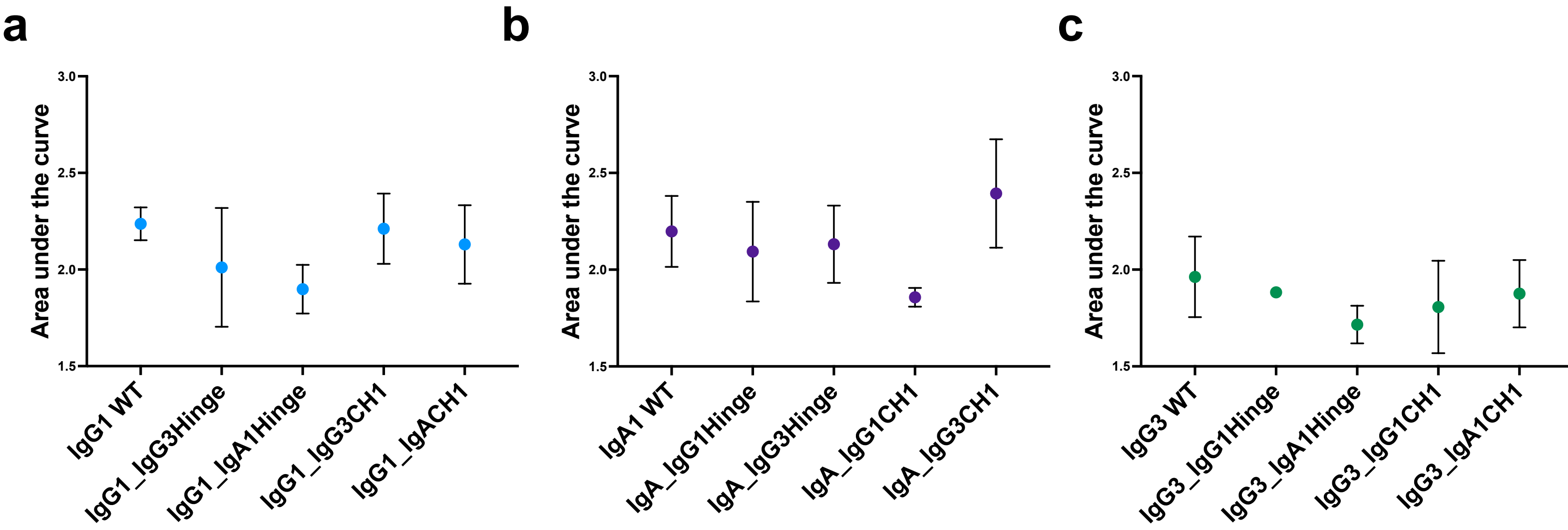
